# Complexity of Genomic Epidemiology of Carbapenem-Resistant *Klebsiella pneumoniae* Isolates in Colombia Urges the Reinforcement of Whole Genome Sequencing-Based Surveillance Programs

**DOI:** 10.1101/2021.06.21.449250

**Authors:** Sandra Yamile Saavedra, Johan Fabian Bernal, Efraín Montilla-Escudero, Stefany Alejandra Arévalo, Diego Andrés Prada, María Fernanda Valencia, Jaime Enrique Moreno, Andrea Melissa Hidalgo, Monica Abrudan, Silvia Argimón, Mihir Kekre, Anthony Underwood, David M Aanensen, Carolina Duarte, Pilar Donado-Godoy, the NIHR Global Health Research Unit on Genomic Surveillance of Antimicrobial Resistance

## Abstract

**Background:** Carbapenem-resistant *Klebsiella pneumoniae* (CRKP) is an emerging public health problem. This study explores the specifics of CRKP epidemiology in Colombia based on whole genome sequencing (WGS) of the National Reference Laboratory at Instituto Nacional de Salud (INS)’s 2013-2017 sample collection.

**Methods:** A total of 425 CRKP isolates from 21 departments were analyzed by HiSeq-X10^®^Illumina high-throughput sequencing. Bioinformatic analysis was performed, primarily using the pipelines developed collaboratively by the National Institute for Health Research Global Health Research Unit (GHRU) on Genomic Surveillance of AMR, and AGROSAVIA.

**Results:** Of the 425 CRKP isolates, 91.5% were carbapenemase-producing strains. The data support a recent expansion and the endemicity of CRKP in Colombia with the circulation of 7 high-risk clones, the most frequent being CG258 (48.39% of isolates). We identified genes encoding carbapenemases *bla*_KPC-3_, *bla*_KPC-2_, *bla*_NDM-1_, *bla*_NDM-9_, *bla*_VIM-2_, *bla*_VIM-4_, and *bla*_VIM-24_, and various mobile genetic elements (MGE). The virulence of CRKP isolates was low, but colibactin (*clb3*) was present in 25.2% of isolates, and a hypervirulent CRKP clone (CG380) was reported for the first time in Colombia. ST258, ST512, and ST4851 were characterized by low levels of diversity in the core genome (ANI> 99.9%).

**Conclusions:** The study outlines complex CRKP epidemiology in Colombia. CG258 expanded clonally and carries specific carbapenemases in specific MGEs, while the other high-risk clones (CG147, CG307, and CG152) present a more diverse complement of carbapenemases. The specifics of the Colombian situation stress the importance of WGS-based surveillance to monitor evolutionary trends of STs, MGE, and resistance and virulence genes.

**summary:** In Colombia, the dissemination of carbapenemases in carbapenem-resistant *Klebsiella pneumoniae* is attributed to horizontal gene transfer and successful circulation of CG258, and, to a lesser extent, other clones such as ST307, ST147, and ST152.

## INTRODUCTION

*Klebsiella pneumoniae* (KP) is a pathogen that causes community and hospital-acquired infections worldwide; it is a threat to public health due to its high levels of antimicrobial resistance [1–3]. Carbapenem-resistant *K. pneumoniae* (CRKP) causes untreatable infections and high mortality [2]. The emergence and global spread of extremely resistant *K. pneumoniae* highlight the need for a greater understanding of resistance epidemiology, for which whole genome sequencing (WGS) methods provide a cost-effective alternative that complements routine methods [4]. Furthermore, the epidemic dynamics of CRKP differ across geographical regions, and it is important to understand local evolutionary and epidemiological variation to establish the most appropriate strategies of surveillance, infection control, and antibiotic stewardship [5, 6].

Colombia’s record of carbapenem use is one of the world’s highest. [7]. According to the last Instituto Nacional de Salud (INS) nationwide report, the use of these antimicrobials exceeds more than 20 times the defined daily dose (DDD) of several countries in Europe and North America [7, 8]. In Colombia, *K. pneumoniae* is the most frequent pathogen found in intensive care units, with resistance to carbapenems reported in up to 15,6 % of isolates [9–12]. The first reports of CRKP associated with KPC-2 were in 2005, and with KPC-3 in 2008 [13, 14]. Colombia is considered KPC endemic [15]. The CRKP clones associated with KPC-type carbapenemases commonly reported in Colombia are CG258, CG307, and CG14/15 [5, 16]. Other carbapenemases identified in CRKP are VIM and NDM-1 (reported since 2011 and 2012 respectively). [12, 17, 18]. The presence of different evolutionary mechanisms in the challenged health care system of Colombia, a medium-income country, may have created the conditions for a significant CRKP epidemic [5].

The present study aimed to extend the scope of previous CRKP reports in Colombia through WGS of CRKP isolates from the 21 most populated departments of Colombia, reported to the National Reference Laboratory (NRL), Instituto Nacional de Salud (INS) between 2013 and 2017. The objective was to increase our understanding of CRKP epidemiology in Colombia to support prevention and control strategies.

## MATERIALS AND METHODS

### Bacterial Isolates

The NRL manages Colombia’s AMR surveillance program and adjusts the criteria for isolate submission according to the epidemiological behavior observed in the country and epidemiological alerts from the Pan American Health Organization (PAHO) (Supplementary Methods).

Between 2013 and 2017, the NRL received 811 clinical isolates of *K. pneumoniae* non-susceptible to carbapenems. After sample selection (Supplementary Materials), 425 isolates remained that met the criteria: 1) from health care-associated infections (HAI) origin; 2) year of isolation between 2013 and 2017; 3) phenotypic confirmation of *K. pneumoniae* bacterial identification by Vitek-2 (bioMérieux); 4) resistance to carbapenems determined by AST disk diffusion using CLSI guidelines for the corresponding year; and 5) genotypic confirmation of carbapenemases in *K. pneumoniae*.

### Whole Genome Sequencing

Genomic DNA was extracted from single-colony cultures using PureLink™ Genomic DNA Mini Kit (Invitrogen, USA). The DNA integrity was checked by electrophoresis in a 1% (w/v) agarose gel, the DNA purity was estimated using NanoDrop^®^ spectrophotometer (Nanodrop Technologies Inc., USA), and the concentration was determined with a Qubit Fluorometer (Invitrogen, USA). Subsequently, 1000 ng of DNA was shipped to the Wellcome Sanger Institute (Hinxton, UK) to sequence using Illumina HiSeq-X10 (San Diego, USA). DNA concentrations were confirmed using AccuBlue Broad Range assay (Biotium, Inc., Fremont, USA), followed by normalization and DNA library construction. DNA was sheared into 450-550 bp fragments using Covaris LC 220 ultrasonicator (Brighton, UK), followed by PCR-based library preparation using Illumina adaptors and 384-indexed tags (NEBNext Ultra II FS DNA library kit). Afterward, size-selection, amplification, purification, and multiplexing were carried out, and libraries were pooled. Pool was quantified and normalized down to 4nM using Biomek NXP workstation for (automated liquid handling; California, USA), Agilent Bioanalyzer 2100 (California, USA), and Roche Lightcycler 480) (Utah, USA), before denaturation and loading on the Illumina platform.

### Quality Control and Bioinformatic Analysis

All genome sequence data were processed using versioned Nextflow workflows and associated Docker containers covering *de novo* assembly, mapping-based SNP phylogeny, MLST assignment, and AMR determinant detection. The workflow structure is summarized in protocols [19]. Major quality control parameters are available in Supplementary Table 1, including file size, number of contigs, genome size, N50. AMR determinants were detected [19], and virulence factor prediction was performed using ARIBA software (github.com/sanger-pathogens/ariba) in conjunction with the VFDB database (github.com/haruosuz/vfdb). MSLT prediction was assigned with ARIBA software using PubMLST databases downloaded on December 20, 2019. The isoforms of Tn*4401* were determined with TETyper (github.com/aesheppard/TETyper) and annotated with Prokka v1.14.5. Additional AMR determinants, virulence factors, and typing (K-locus, wzi, and O-locus) were detected and species predicted using Kleborate, and Kaptive as implemented on the PathogenWatch platform (https://pathogen.watch) [20, 21].

SNP variants were called relative to a reference genome (*K. pneumoniae* strain K2044 (NCBI RefSeq NZ_CP026011.1)) and a maximum likelihood phylogeny inferred from pseudogenome alignment of the variants using IQ_TREE with GTR+G model and 1000 bootstrap replicates [19, 22]. High-quality views of trees and data were rendered using the ggtree R package (bioconductor.org/packages/ggtree). For clustering, clonal groups were defined using core genome SNP analysis using Prokka v1.14.5 gene annotation and a 70-90% ANI between isolates.

Raw sequence data were deposited at the European Nucleotide Archive under accession <u>PRJEB29742</u> and visualized with Microreact (https://microreact.org/project/vcsgT8Ic4/66d1f105) [23].

## RESULTS AND DISCUSSION

### Geographic Spread of High-Risk Clones in Colombia

The 425 CRKP isolates analyzed in this study originated from 21 departments of Colombia, representing 90% of the total population (Figure 1a). These isolates belonged to 129 institutions (114 hospitals and 15 microbiology laboratories). The level of medical services was determined for 102 institutions, of which 68, 19 and 15 provided high, medium-high, and medium levels respectively.

**Figure 1.**
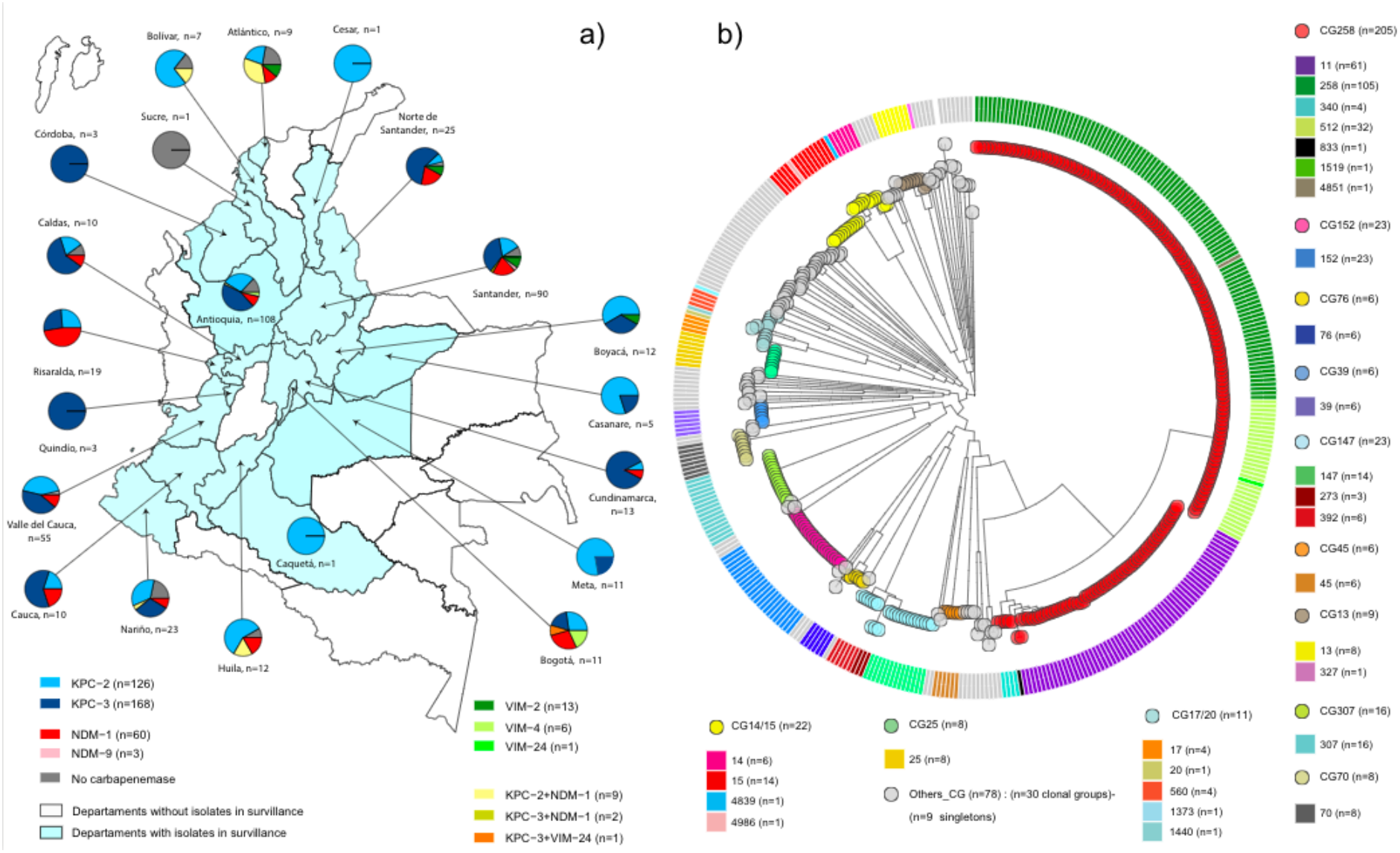
Geographical distribution of mechanisms of resistance to carbapenems in *Klebsiella pneumoniae* and core genome tree in Colombia, 2013-2017. a) Map depicting the 21 departments that submitted isolates for surveillance. The isolates were classified according to the mechanism of resistance to carbapenem by color (see key). Pie chart shows the proportions of different mechanisms of resistance to carbapenems in each department. b) Circular representation of the transformed phylogenetic tree showing the genetic relationships among 421 Colombian *K. pneumoniae* CRKP isolates. The phylogenetic tree was built using 421/425 isolates; 4 were excluded from the analysis because they presented a high number of N in the multiple alignments. The 421 isolates were classified into 80 sequence types (STs) grouped in 42 CG and 9 singletons. Due to the number of different CGs, for the graph, CGs with ≥6 isolates were selected for easy viewing on the tree: CG258, CG13, CG14/15, CG17/20, CG25, CG39, CG45, CG70, CG76, CG147, CG152, and CG307. The remaining CG and singletons were classified as other-CG.

Of all isolates, 93.9% (n = 399) were multidrug-resistant (MDR), and 40.5% (n=172) were resistant to at least one antibiotic from all classes tested (Supplementary Table 2).

A total of 80 sequence types (STs) were identified; 42 STs were reported for the first time in Colombia, including 10 novel STs: STs 4838-4841, 4851, 4852, 4984–4987 (Supplementary Table 2). The STs were classified into 42 clonal groups (CGs) and 9 singletons (Figure 1b). The most frequent CGs were: CG258, CG147, CG152, CG14/15, CG307, CG17/20, and CG13 (Figure 1b and Supplementary Table 3). These include known epidemic (“high-risk”) STs that are over-represented in the global public genome collections. Of the 7 most frequent CGs of this study, 6 had been previously reported in Colombia [5, 15, 16, 21, 24–29]. To the best of our knowledge, this is the first report of ST152 in the Americas.

CG258, the main CG identified in this study (48.9%, n=208), presented the broadest geographic distribution (18 departments). CG17/20 (2.6%, n=11) was reported in 7 departments. CG147 (5.4%, n=23) and CG 14/15 (5.2%, n=22) were present in 6 departments each, and CG152 (5.4%, n=23) in 5 departments. Three geographically close departments, Antioquia, Santander, and Valle del Cauca (60%, n=255) reported the simultaneous presence of the 7 main CGs.

All but one of the 7 main CGs showed increased and recent dissemination compared to previous reports in Colombia. Of CG258, ST11 was reported in 10 departments in this study, contrasting with previous reports, in which it was identified in only 2 departments [16, 30]. ST512 which was previously only detected in Medellin, is now reported in 4 departments [5, 16]. CG147 and CG307, which had been sporadically reported in Antioquia, were reported in 11 and 6 departments, respectively [16, 26, 29, 31]. CG152 is reported for the first time in Colombia in 5 departments (Supplementary Figure 1a). In this study, the prevalence of CG14/15 (5.2%) is lower compared to previous reports, of 30% between 2005 and 2007 and 8.3% between 2012 and 2014 [5]. This study represents the broadest geographical analysis of CRKP in Colombia and confirms their endemic status.

Most of the STs found in Colombia in this study have been reported globally. ST258, the dominant high-risk ST worldwide, was first reported in Colombia in 2006 following global clonal propagation [15, 21, 24, 25]. CG258 (ST258, ST11, ST512) has been identified as responsible for 68% of 112 *K. pneumoniae* outbreaks worldwide, reported between 2005 and 2016 [32]. The percentage of CRKP that belongs to CG258 in this study is lower than those reported in the USA (70%) by the Centers for Disease Control and Prevention (CDC), or in Israel (90%) [33–35]. ST11 is a prevalent clone in Asia, while its spread in Latin America had been only reported in Brazil [36–39]. ST512 is endemic in Italy, Israel, and Greece, ST147 in India, Italy, Tunisia, and Greece, and CG307 in Italy and the United States [16, 29, 31, 40–42]. Since differences in fitness, epidemicity, and host-related factors between CGs have been reported, molecular identification of specific high-risk clones in a region is of the utmost importance to implement appropriate infection control programs [32].

### Detection and Distribution of Antimicrobial Resistance Mechanisms and Mobile Genetic Elements

Carbapenemase genes were identified in 389 CRKP genomes; 36 isolates were non-carbapenemase producers.

#### *bla*_KPC_

Among carbapenemase, *bla*_KPC_ was identified in 294 isolates, and *bla*_NDM_ and *bla*_VIM_ in 63 and 20, respectively. Co-production of *bla*_KPC+NDM_ was found in 11 isolates and co-production of *bla*_KPC+VIM_ in 1 isolate (Figure 1a). These findings confirm that, unlike in other regions of the world, *bla*_KPC_ continues to dominate in Colombia, and *bla*_OXA-48-like_ does not play a role in carbapenem resistance. They also indicate a recent expansion of *bla*_NDM_ in Colombia.

We found two variants of *bla*_KPC_ – *bla*_KPC-3_ (39%, n=168) and *bla*_KPC-2_ (30%, n=126) – across 64 different STs in multiple departments (Figure 1a). Additionally, a wide variety of mobile genetic elements with *bla*_KPC_ were identified. The most common plasmid replicons in *bla*_KPC_-positive isolates were IncFIB(K) (86.7%), IncFII(K) (56.8%), ColRNAI (53.1%), IncR (39.1%), and IncFII (32.3%) (Table 1, Figure 2c). From the KPC-carrying isolates, 68% showed the presence of the Tn*4401* transposon. In the isolates carrying *bla*_KPC-2_, the gene was located in three Tn*4401* isoforms and in non-Tn*4401* elements (NTEKPC) (n=52) (Table 1, Figure 2d). In contrast, *bla*_KPC-3_ was localized in Tn*4401*a in ST512 and ST1519 isolates and in Tn*4401*b in 82.5% of ST258. The clonal lineage ST258/512 accounts for 47% of all KPC genes reported in this study, while this percentage is >70% in the EuSCAPE sample collection [43]. The variant *bla*_KPC-2_ is associated with a greater variety of mobile genetic elements than in previous reports and is found in different CGs, including CG307, CG14/15, and CG147 (Table 1), suggesting that its propagation is associated with successful Horizontal Gene Transfer (HGT) [5, 17]. In contrast, *bla*_KPC-3_ is correlated with the presence of two types of Tn*4401* elements in the clonal lineage ST258/512, suggesting dissemination by clonal expansion. These observations confirm and extend the suggested compartmentalization of the genes encoding for the two KPC variants in Colombia by Rojas *et al*. [5]. This horizontal transfer and promiscuity of KPC-2-carrying mobile genetic elements might have appeared in Colombia by selective pressure due to increased use of carbapenems [7].

**Figure 2.**
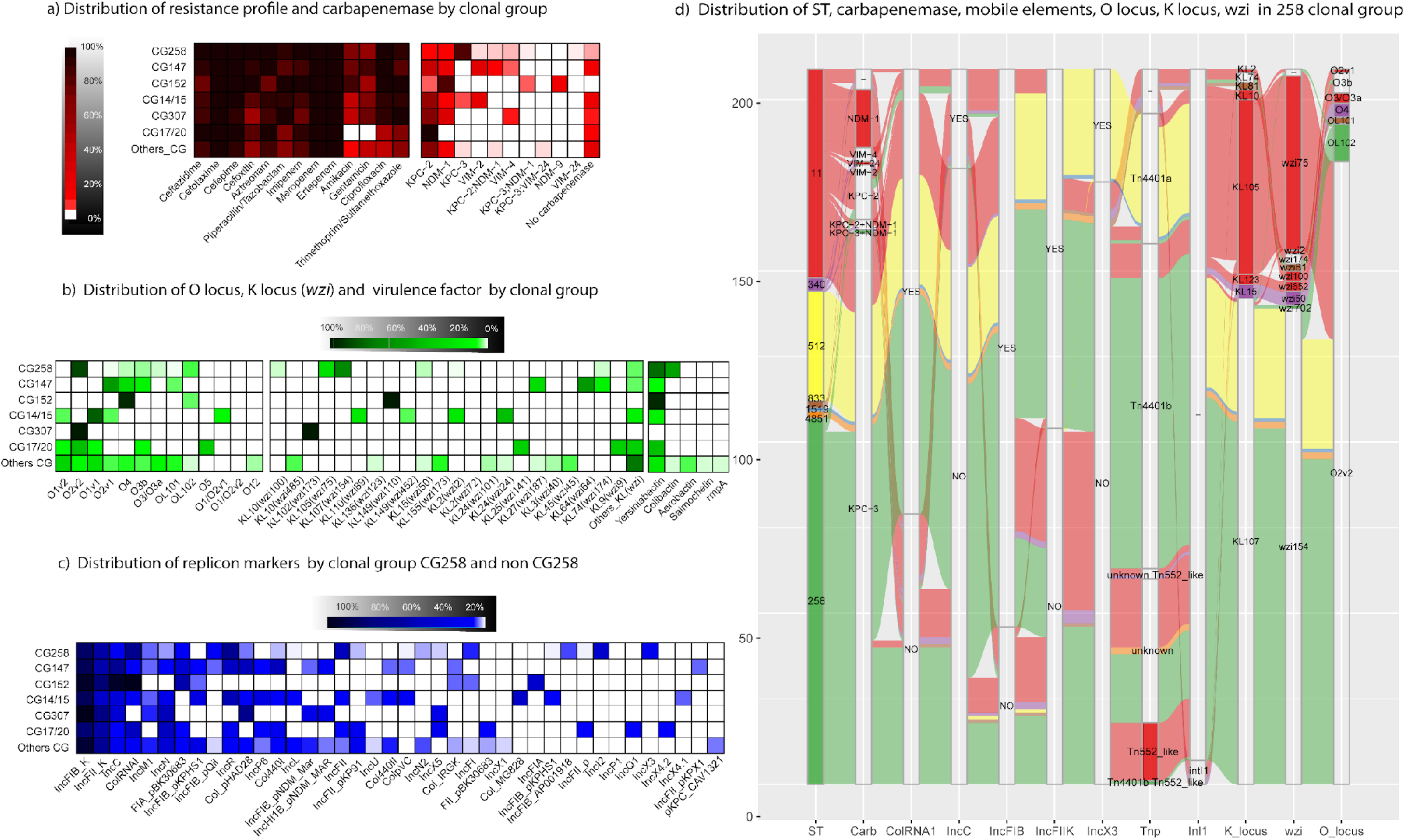
Clonal group heat maps, virulence factors, resistance profile, carbapenemases, replicons, and alluvial of the CG258. The heatmap analyzes different clonal groups (each row of the heatmap represents a clonal group). a) Heatmap displaying the distribution of antimicrobials and the mechanism of resistance to carbapenems (y-axis) in the clonal group. b) Heatmap displaying the distribution of O locus, K locus (*wzi*), and virulence factors (y-axis) found in the clonal group. c) Heatmap displaying the distribution of the replicon type (y-axis) found in the clonal group. d) Alluvial diagram showing the “flow” of the presence of the most important transposable elements associated with carbapenemase and ST of CG258.

**Table 1.**
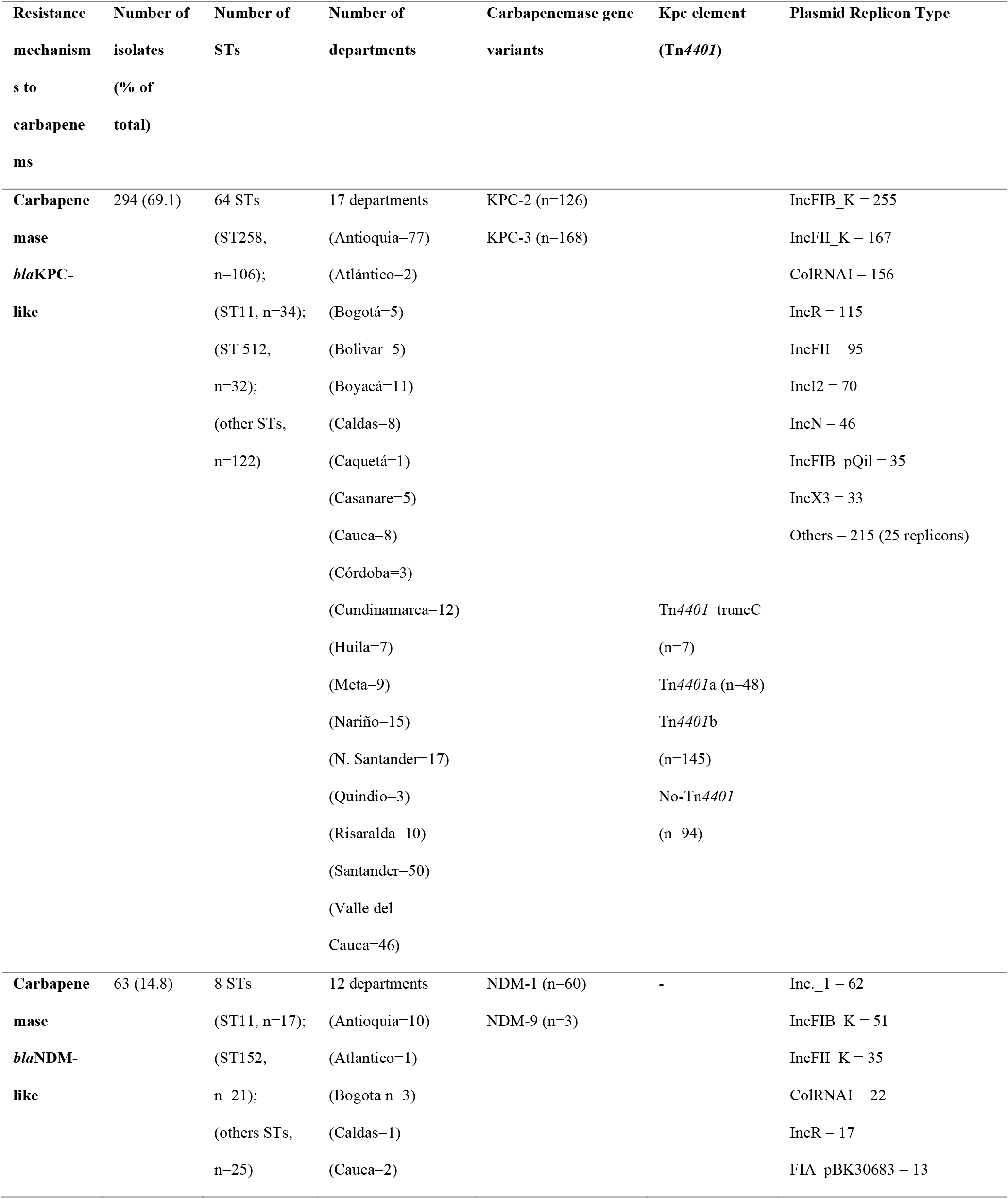
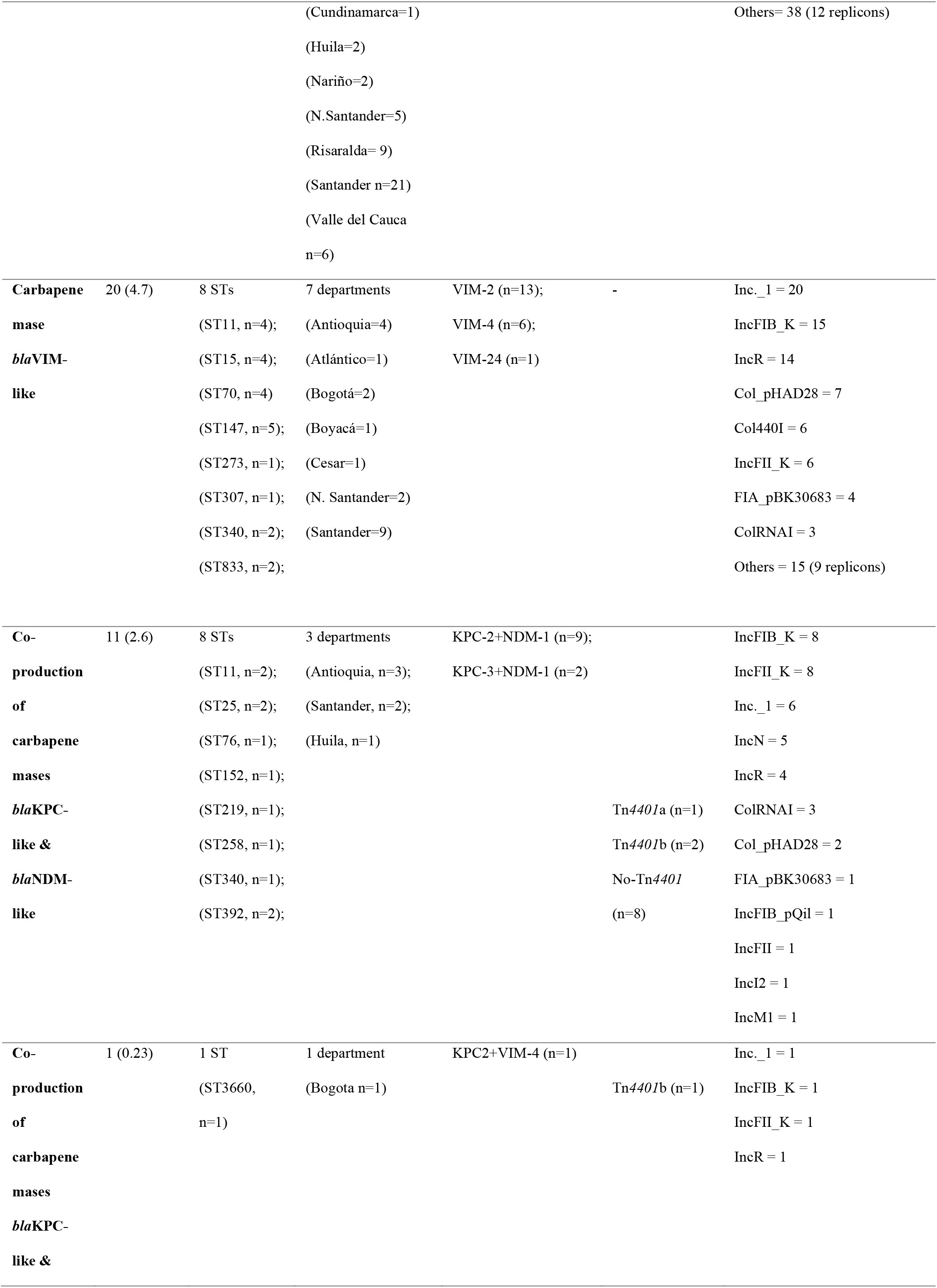
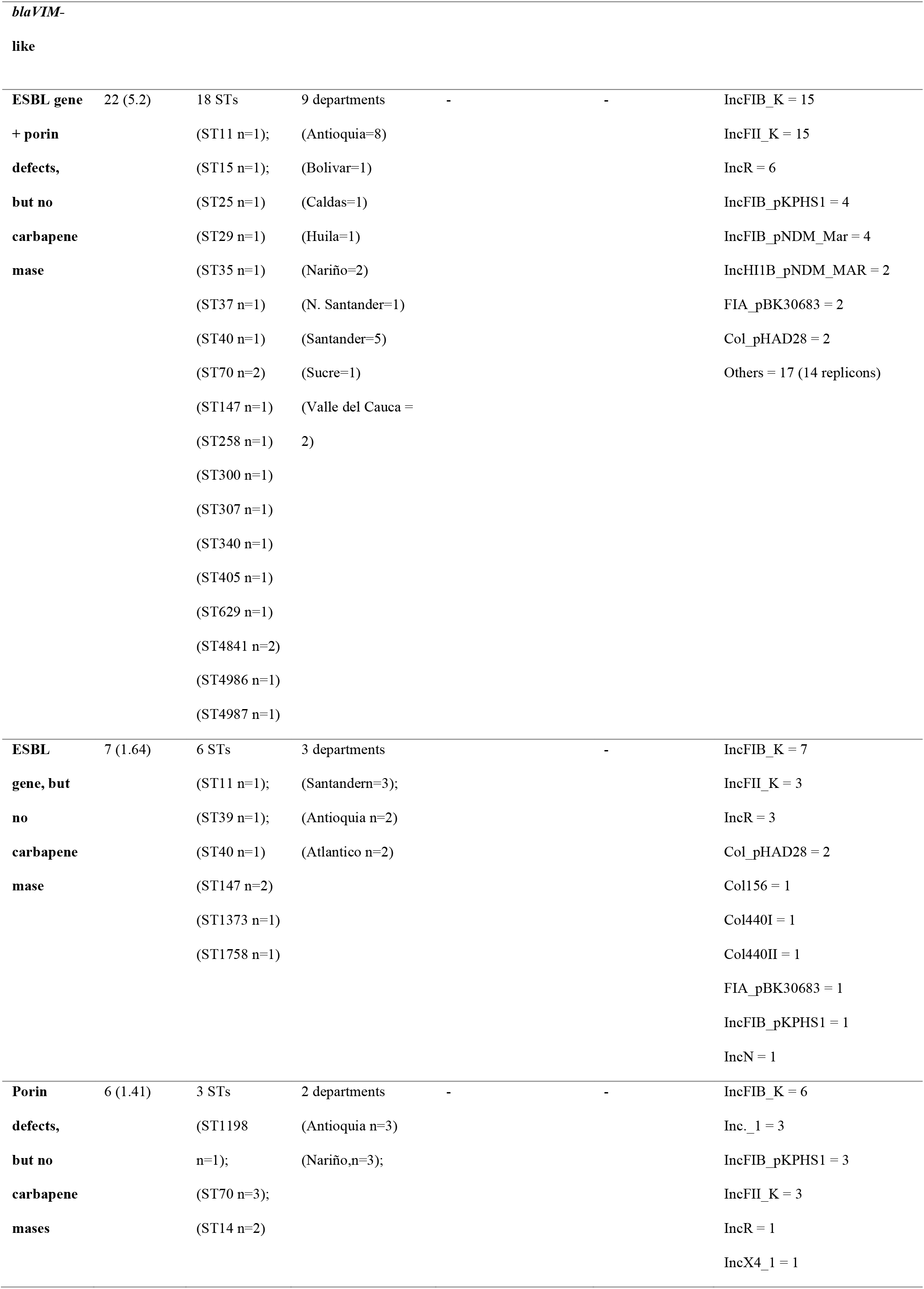
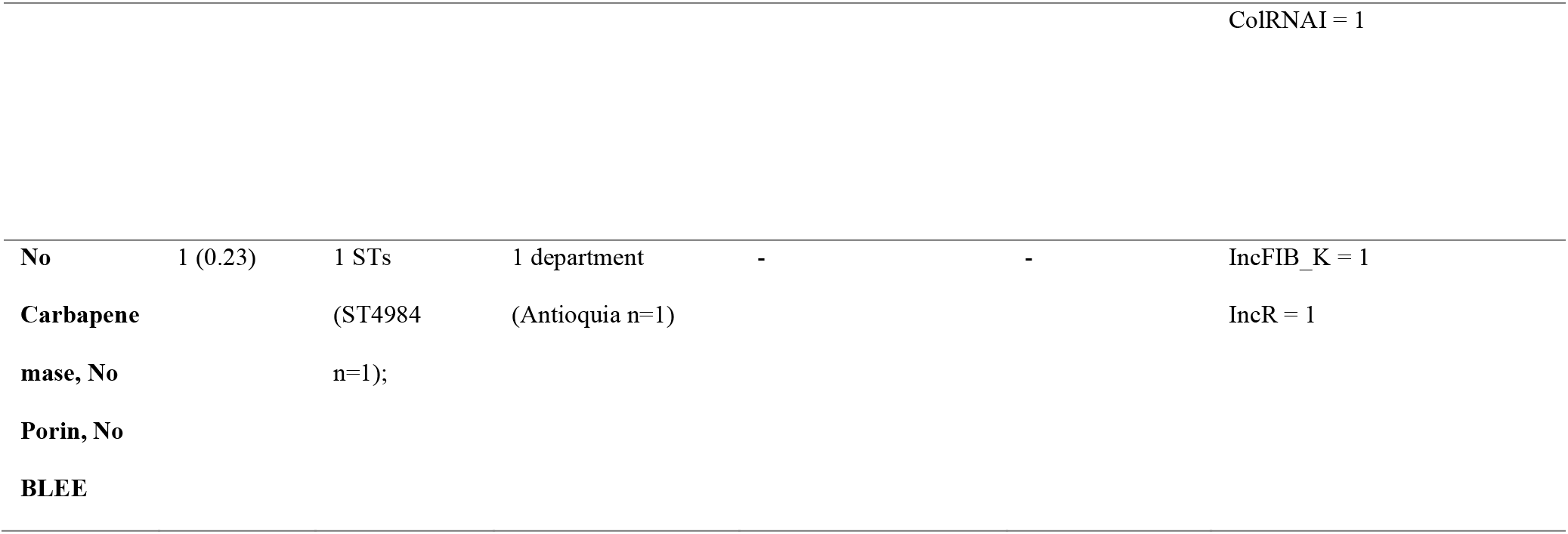
Characteristics of carbapenem-resistant *K. pneumoniae* isolates.

#### *bla*_NDM_

Two variants of *bla*_NDM_ were found: *bla*_NDM-1_ (n=60) and *bla*_NDM-9_ (n=3). This is the first report of *bla*_NDM-9_ in the Americas. Also, it is the first time that the dissemination of *bla*_NDM-1_ is reported in 12 departments in Colombia (Figure 1a). The two variants were associated mainly with ST152 (33%) and ST11 (27%) (Supplementary Figure 1b). An IncC plasmid was carried by 98.4% of *bla*_NDM_ isolates. Co-location of both determinants (IncC-*bla*_NDM_) in the same contig could be confirmed in 15.9% of isolates, using Artemis (Supplementary Figure 2). The IncC replicon is common in *bla*_NDM_ isolates worldwide and has been previously reported in Colombia [28, 44]. Other isolates showed incomplete replicons due to short read sequencing limitations. The genetic context was identical in all isolates, including variants *bla*_NDM-1_ and *bla*_NDM-9_ (Supplementary Figure 2). A Tn*Ec*-like transposon from Tn*3* family was identified in all genomic contexts with NDM genes (within 3093 bp) [28]. We conclude that the presence of NDM is associated with the expansion of the clones ST152 and ST11 and horizontal transfer of NDM-1 via the vehicle of an IncC plasmid.

#### *bla*_VIM_

Three *bla*_VIM_ variants – *bla*_VIM-2_ (n=13), *bla*_VIM-4_ (n=6) and *bla*_VIM-24_ (n=1) – were reported in 20 isolates from 7 departments, belonging to 8 STs, mainly ST147 (23.8%, n=5), ST11 (19%, n=4), and ST15 (19%, n=4). The 3 *bla*_VIM_ variants were found within 2 different class 1 integrons, known to be responsible for the mobilization of VIM genes (Supplementary Figure 2) [42, 45]. Our results contrast with those reported globally, where *bla*_VIM-1_ is the most common variant in VIM-carrying CRKP [42]. The observation that *bla*_VIM-2_ is the most common allele of VIM in *P. aeruginosa* in Colombia suggests that HGT may have been responsible for the intra-species transfer of the gene [45]. Further work will be carried out to investigate this hypothesis and the potential directionality of the transfer.

#### Co-productions

The 11 isolates with co-production of *bla*_KPC+NDM_ came from 3 departments and were associated with 8 STs. In the isolates co-harboring *bla*_KPC-3+NDM-1_, it was observed that *bla*_KPC-3_ incorporated with Tn*4401*a exhibited a genetic environment, including [trpR (Tn3)-ISPsy42(Tn3)-ISKpn7(IS21)-blaKPC-3-ISKpn6(IS1182)-aacA4], and that the vehicle for *bla*_NDM-1_ was similar (TnEc-like). Co-producers of *bla*_KPC-2+NDM-1_ contained the same *bla*_NDM-1_ platform (TnEc-like), but the *bla*_KPC-2_ genetic environment revealed a diversity of genomic contexts. In the isolate co-producers of *bla*_KPC-3+VIM-24_, the *bla*_VIM-24_ gene was found associated with class-1 integron in IncC, and *bla*_KPC-3_ exhibited Tn*4401*b complemented with *Tn2* transposases (Supplementary Figure 2). CRKPs with co-production of at least 2 genes have been previously reported worldwide, including *bla*_KPC_ and *bla*_VIM_ in Italy, *bla*_KPC-2+NDM-1_ in China and Brazil, and *bla*_NDM-1+OXA-181_ in Singapore [46]. The 3 co-productions reported in this study have been previously sporadically reported in Colombia [11, 12, 28], but this is the first report of the full replicon. Monitoring the dynamics of CRKPs with co-productions in the surveillance program is essential because of the potential risks they present, and since the way they emerge and whether they are only transient or have transmission capacity remain uncertain [43, 46].

#### CRKP isolates not harboring carbapenemases

In 36 CRKP isolates not harboring a carbapenemase gene in their genomes, the resistance to carbapenems could be attributed to the presence of extended spectrum β-lactamase (ESBL) genes (86%, n = 31), especially *bla*_CTX-M-15_ (22/31); in 24 of the 31 isolates, these ESBL genes were in combination with porin truncations (Supplementary Table 2).

Porin truncations or mutations were found in 52% of all isolates. These truncations result in a nonfunctional pore that acts in concert to lower carbapenem concentrations, limiting antibiotic influx [47]. In one isolate, no resistance mechanisms were identified (Supplementary Table 3).

The enzymatic hydrolysis of carbapenems by carbapenemases was identified as the primary mechanism of carbapenem resistance, and the vast majority of carbapenemase producer isolates (91%, n=385) were resistant simultaneously to all screening carbapenems (IPM, MEM, ETP) (Supplementary Figure 3). Possible explanations include the preferential use of carbapenems in hospitals for the treatment of serious infections caused by carbapenem resistance isolates, as well as the circulation of *bla*_KPC-2_ and *bla*_KPC-3_-harboring isolates promoting the progression of carbapenem resistance in Colombia, which is a KPC endemic country.

Additional AMR determinants to aminoglycosides (n=390), fluoroquinolones (n=378), phenicols (n=317), ESBL and/or AmpC (n=248), and tetracyclines (n=219) were observed in the majority of isolates (Supplementary Table 2).

Our results outline the increasing complexity of the CRKP population, resistance mechanisms, and lineages in Colombia, which will require adaptations of the conventional infection control measures that were successful in areas where the objective was to curb the expansion of a single dominant clone.

### Detection of Virulence Mechanisms and Serotypes

We observed a low prevalence of acquired virulence mechanisms among CRKP isolates in this study, which is in line with global trends [48]. However, we additionally observed an over-representation of colibactin genes among the isolates and the first observation in Colombia of hyper-virulent CRKP. The CG258 isolates exhibited different types of K-locus (KL), *wzi*, and O-locus (OL). The most common profile was KL107-*wzi*154:O2v2 (STs 258, 512, 1519, and 4851), present in 62.5% of isolates (Figure 2a, Figure 2b, Supplementary Table 3) [21, 40, 49, 50]. The profile KL105-*wzi*75:O2v2, detected in 24% of isolates, is unusual for ST11 [21]. In China, where ST11 is endemic, the dominant KLs are KL47 and KL64, but the occurrence of capsule switching has been suggested in the case of this particular ST [37, 51]. In CG258, 76.9% of isolates were carriers of yersiniabactin (*ybt*), mainly of the *ybt*17 (105 isolates of ST258 and 2 of ST4851) and *ybt*10 (50 isolates of ST11 and 1 of ST258) lineages (Figure 2a, Figure 2b, Supplementary Table 3). Details for other CGs are shown in Figure 2a and Supplementary Table 3.

It is remarkable to note that isolates of ybt17 / colibactin (*clb3*) lineages represented 25.2% of all isolates, while the spread of *clb3* among CRKP is usually around 10% [52]. However, most isolates *ybt*17+*clb*3+ST258 have been reported in the literature as not being colibactin producers, due to a deletion in *clb*J/*clb*K [50, 52].

Only one isolate (RB 877) from this study exhibited determinants associated with hypervirulence, such as *ybt*16, aerobactin (*iuc*2), salmochelin (*iro*2), and *rmpA_3* (KpVP2). This ST380 isolate was recovered in Antioquia in 2013 and is a producer of KPC-2, with the profile KL2-*wzi*203:O1v1 (Table 1, Supplementary Table 2). CG380 has been classified as a hypervirulent clone [48]. This is the first time a hypervirulent CRKP is reported in Colombia.

The two cases of virulence elements reported in this study illustrate the risk of convergence of resistance and virulence, either through the acquisition of virulence plasmid(s) by CRKP or through the acquisition of resistance mobile genetic element(s) by virulent clone. The latter appears to be more difficult [52]. This emphasizes the need for genomic surveillance programs to include both resistance and virulence locus information.

### Genetic Diversity Among CRKP Isolates

The phylogenetic analysis by SNP reflected the MLST distribution into STs as described above. Additionally, it highlights the genomic diversity within each ST. Of interest, we observed ST11, ST147, ST152, and ST307 with low nucleotide divergence (0.05 - 0.47%), suggesting clonality (Figure 1b). However, a deeper analysis by ST showed distinct clades within each ST (Supplementary Figure 4).

Among the 208 isolates of CG258 (48.9% of the total isolates) grouped into 7 STs (Figure 1b and 2b), the ST258, ST512, and ST4851 were characterized by low levels of phylogenetic diversity (ANI> 99.9%). This indicates a recent and common evolutionary ancestor that supports their grouping. The remaining STs of CG258 showed greater diversity (ANI 99.7-99.8%) among pairs. ST11 isolates carried a wide variety of carbapenemases (KPC, NDM, VIM). To our knowledge, this is the first report of *bla*_NDM+VIM_-carrying ST11 isolates in Colombia, although they have been reported worldwide [42, 44, 53]. There were several distinct clades of ST11, with different serotypes, resistance genes, and plasmid replicons (Supplementary Figure 4). This opens the possibility of multiple introductions or emergence of ST11 into Colombia and should be further investigated by contextualizing the genomes with global collections.

The ST147 showed several clades with differences in carbapenemase genes, related to Inc-type elements, discarding the clonality of the group (Supplementary Figure 4). Similarly, ST307 showed different clades some with *bla*_KPC_, *bla*_NDM_, or *bla*_VIM_, alongside IncFIB mobile elements and ESBL genes. The apparent clonality may come from the acquisition of carbapenemase genes in multiple events, related to mobile elements. This is in accordance with ST147 found related to Inc-type plasmids, and ST307 to *bla*OXA48-like plasmid genes, both related to carbapenemase genes HGT [43]. ST152 was found as an emerging clone with a genomic profile characterized by *bla*_NDM_, several Inctype mobile elements, and hypervirulence provided by the presence of yersiniabactin, similar to that reported in Saudi Arabia and other countries as a hypervirulent ST of importance in urinary tract infections [54]. All previous analyses enhances the importance of WGS for AMR surveillance, as differential findings between STs and genes reveal non-clonality that may be translated into differential treatment of CRKP infections.

Among this study’s limitations were: having only voluntary surveillance isolates at the national level; and limited data of clinical and epidemiological information for a more exhaustive investigation. Nevertheless, the information obtained is essential for national surveillance and provides insights into pathogen genetics for planning effective surveillance.

## CONCLUDING REMARKS

This study allowed a detailed description of CRKP epidemiology in Colombia that should be taken into account when establishing efficient WGS-based surveillance programs in the country.

There are 7 predominant high-risk STs in Colombia. In addition to the previously described CG258 with KPC, we showed a complex, recent and significant expansion of multiple clones of interest and carbapenemases, urging reinforcement of the CRKP genomic surveillance process.

The dissemination of carbapenemase genes is based on a complex combination of mechanisms. KPC-3 occurs by the expansion of CG258, mainly ST258 and ST512 carrying an IncFII replicon plasmid with the insertion of transposon Tn*4401*-like. The spread of KPC-2 and VIM genes was associated with the horizontal transfer of different plasmid replicons and mobile elements. NDM-1 dissemination was associated with the expansion of clones CG152 and ST11 (CG258) and possible HGT related to IncC. The high proportion of HTG mechanisms in disseminating carbapenem resistance genes could have resulted from higher selective pressure due to overuse of antibiotics in a challenged health care system. This result shows the importance of including mobile genetic elements in the CRKP surveillance program in Colombia.

The hypervirulent strain report and the over-representation of the colibactin virulence factor urge close monitoring of the potential convergence of virulence and resistance of CRKP in Colombia.

## Supporting information

Table S1. Detailed quality control (QC) results for all sequenced isolates submitted as K.pneumoniae

Table S2. Demographic, phenotypic and genotypic characteristics of 425 isolates of carbapenem-resistant Klebsiella pneumoniae

Table S3 Correlation CG, ST, Department, Mechanism resistance to carbapenem, K - locus, wzi, O-locus, virulence factor in the Clonal Group

## ABBREVIATIONS

CRKP: Carbapenem resistant *Klebsiella pneumoniae*
KPC: *Klebsiella pneumoniae* carbapenemase

## ACKNOWLEDGMENTS

Members of the NIHR Global Health Research Unit on Genomic Surveillance of Antimicrobial Resistance: Khalil Abudahab, Harry Harste, Dawn Muddyman, Ben Taylor, Nicole Wheeler, and Sophia David of the Centre for Genomic Pathogen Surveillance, Big Data Institute, University of Oxford, Old Road Campus, Oxford, United Kingdom and Wellcome Genome Campus, Hinxton, UK; Alejandra Arevalo, and Erik C. D. Osma Castro of the Colombian Integrated Program for Antimicrobial Resistance Surveillance – Coipars, CI Tibaitatá, Corporación Colombiana de Investigación Agropecuaria (AGROSAVIA), Tibaitatá – Mosquera, Cundinamarca, Colombia; K. L. Ravikumar, Geetha Nagaraj, Varun Shamanna, Vandana Govindan, Akshata Prabhu, D. Sravani, M. R. Shincy, Steffimole Rose, and Ravishankar K.N of the Central Research Laboratory, Kempegowda Institute of Medical Sciences, Bengaluru, India; Iruka N Okeke, Anderson O. Oaikhena, Ayorinde O. Afolayan, Jolaade J Ajiboye, and Erkison Ewomazino Odih of the Department of Pharmaceutical Microbiology, Faculty of Pharmacy, University of Ibadan, Oyo State, Nigeria; Celia Carlos, Marietta L. Lagrada, Polle Krystle V. Macaranas, Agnettah M. Olorosa, June M. Gayeta, and Elmer M. Herrera of the Antimicrobial Resistance Surveillance Reference Laboratory, Research Institute for Tropical Medicine, Muntinlupa, the Philippines; Ali Molloy, alimolloy.com; John Stelling, The Brigham and Women’s Hospital; and Carolin Vegvari, Imperial College London.

The authors acknowledge all public health laboratories from the National Reference Laboratory (NRL) for providing the isolates CRKP obtained under the previous National Surveillance by the Laboratory of Antimicrobial Resistance Surveillance Network.

We acknowledge Angela Sofía García, MSc, part of the Colombian Genomic Unit of AGROSAVIA, for the valuable suggestions on the text to enrich the content of this article.

