## Supplementary Information for "Complexity of Genomic Epidemiology of Carbapenem-Resistant *Klebsiella pneumoniae* Isolates in Colombia Urges the Reinforcement of Whole Genome Sequencing-Based Surveillance Programs"

[a] Members of the NIHR Global Health Research Unit on Genomic Surveillance of Antimicrobial Resistance are listed in the Acknowledgments.

** Corresponding author

### SUPPLEMENTARY METHODS

**Description of Colombian AMR Surveillance: Antimicrobial Resistance Program in Health Care-Associated Infections (*Infecciones Asociadas a la Atención en Salud* IAAS)**

The AMR surveillance by the laboratory in healthcare-associated infections (HAI) modifies the criteria for isolates submission according to the epidemiological behavior observed in the country and the epidemiological alerts from the Pan American Health Organization (PAHO) / World Health Organization (WHO).

For 2013, isolates resistant to at least one 3rd-generation cephalosporin with resistance or decreased susceptibility to at least one carbapenem were received following the guidelines 057 of 2012 (https://www.ins.gov.co/Normatividad/Circulares/CIRCULAR%200057%20DE%202012.pdf#search=circular%20057), and 043 of 2013 (https://www.ins.gov.co/Normatividad/Circulares/CIRCULAR%20EXTERNA%200021%20DE%202014.pdf#search=circular%20021%20de%202014).

For 2014 and 2015, the referral was carried out according to guideline 021 of 2014, meaning: i) isolates resistant to at least one 3rd-generation cephalosporin and with resistance or decreased susceptibility to at least one carbapenem; ii) positive results to phenotypic ethylenediaminetetraacetic acid (EDTA)/sodium mercaptoacetic acid (SMA) test; and iii) the first isolation of carbapenemase-producing enterobacteria from each institution after screening with modified Hodge test (MHT) and synergy with boronic acid (APB) disc test.

In 2016, a delivery criteria flowchart was established, and this is in force to date (https://www.ins.gov.co/buscador-eventos/Informacin%20de%20laboratorio/Criterios-para-env%C3%ADo-de-aislamientos-bacterianos-y-levaduras-del-g%C3%A9nero-C%C3%A1ndida-en-IAAS.pdf ). In general, the same criteria for the submission of isolates established for 2014-2015 were maintained, while isolates with a negative result to EDTA / SMA and APB test were included.

**Sample Selection**

From 2013 to 2017, the NRL received 811 clinical isolates of *K. pneumoniae* non-susceptible to carbapenems. Upon reception, bacterial identification was performed using ≈ 2 (bioMérieux), antimicrobial susceptibility was determined by disk diffusion, and results were interpreted using CLSI guidelines for the corresponding year. Phenotypic detection of carbapenemases was performed with Hodge test, EDTA test and acid 3-aminophenylboronic test (APB), while molecular detection was carried out using PCR for any positive CRKP, following Ovalle et al [1].

Out of the 811 confirmed as carbapenems resistance isolates, 557 were selected to be sequenced, the isolates were recovered on MacConkey agar, and their species and susceptibility to carbapenems were re-confirmed with VITEK 2 (bioMérieux), following the interpretation breakpoints from CLSI 2020. Of these, 132 samples were subsequently excluded, either because the isolates became nonviable, were identified as a species other than *K. pneumoniae,* or were not confirmed as CRKP.

#### Supplementary Tables

**Supplementary Table 1.** Detailed quality control (QC) results for all sequenced isolates submitted as *Klebsiella pneumoniae*.

[Provided as Excel spreadsheet]

**Supplementary Table 2.** Demographic, phenotypic and genotypic characteristics of 425 isolates of carbapenem-resistant *Klebsiella pneumoniae*.

[Provided as Excel spreadsheet]

**Supplementary Table 3.** Correlation CG_ST_Department Mechanism resistance to carbapenem K-locus and *wzi* O-locus virulence factor in the clonal group and singletons.

[Provided as Excel spreadsheet]

#### Supplementary Figures

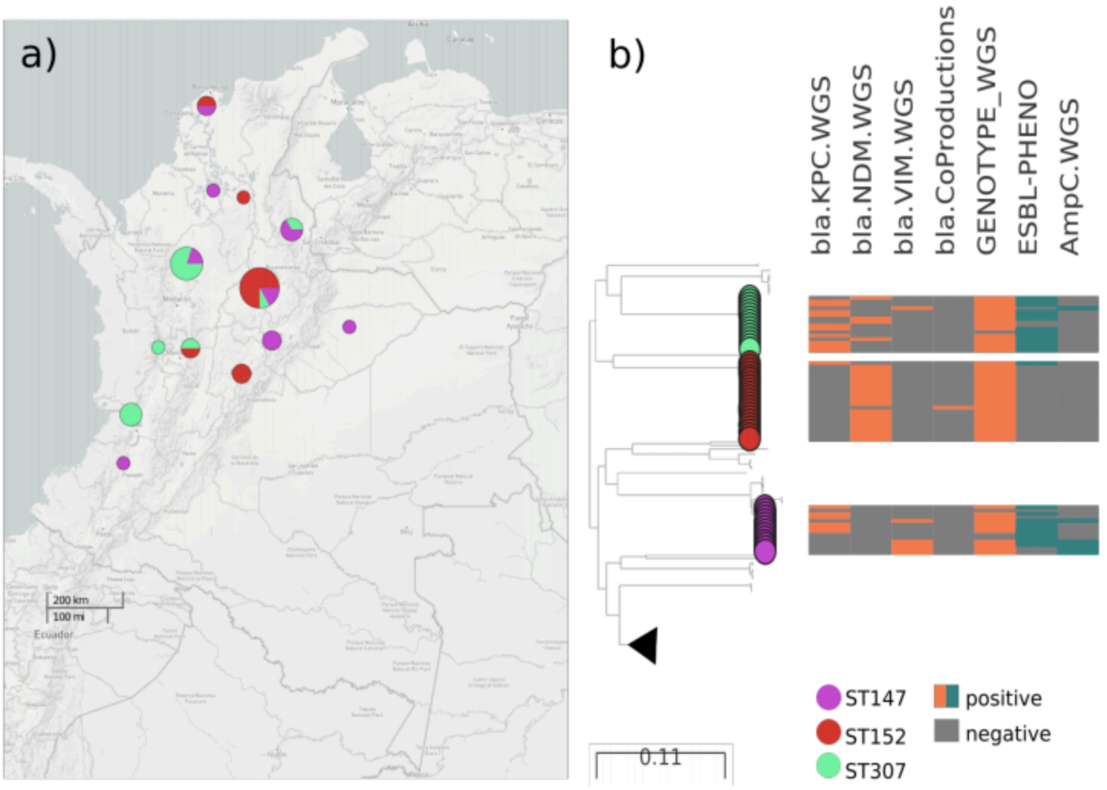


**Supplementary Figure 1.** Geographic and phylogenetic distribution of ST147, ST152, and ST307.

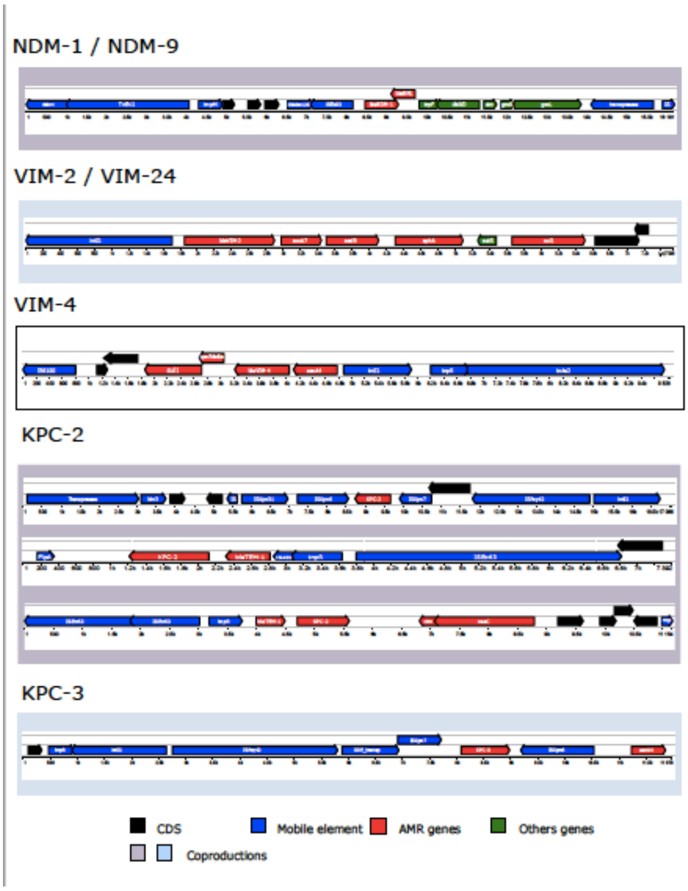


**Supplementary Figure 2.** Carbapenemase genetic environments. (Available as pdf.)

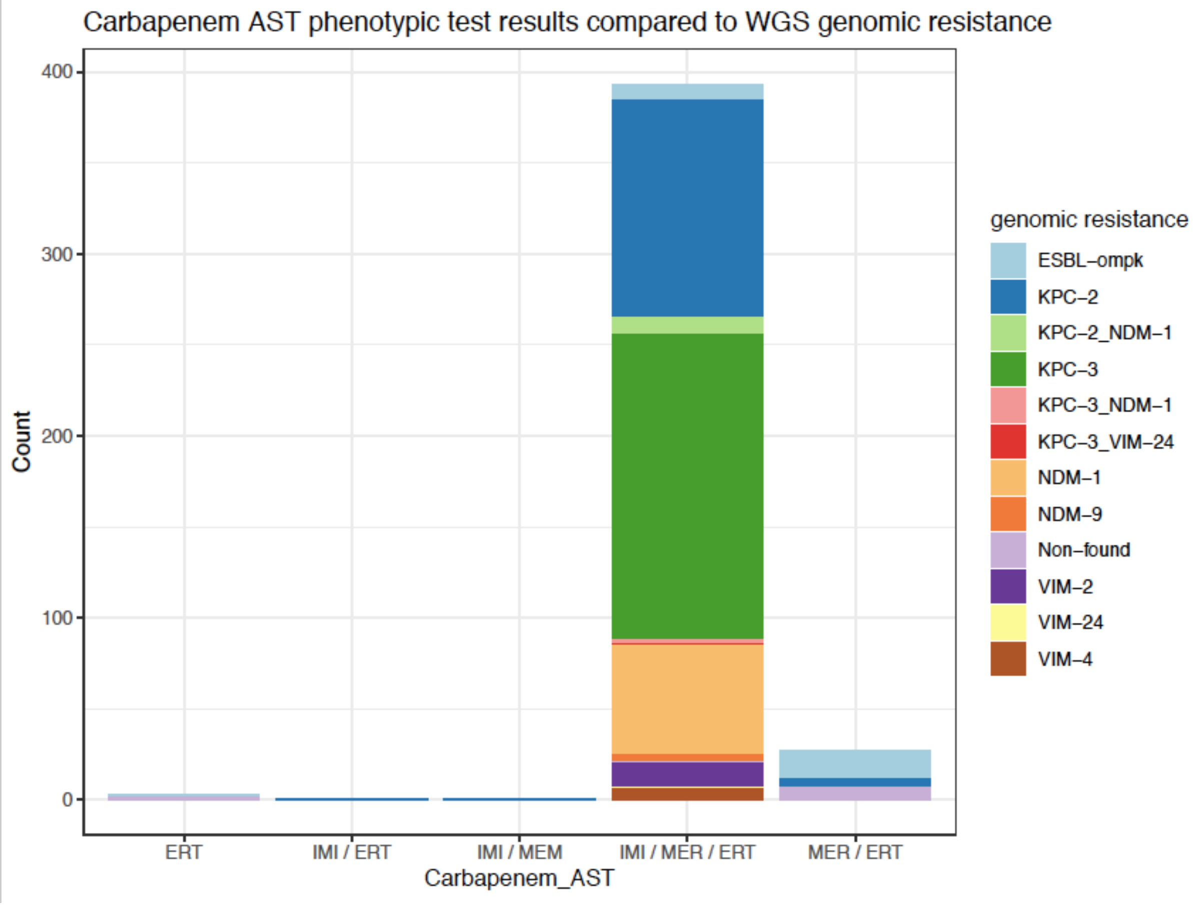


**Supplementary Figure 3.** Carbapenem AST phenotypic test results compared to WGS genomic resistance. (Available as pdf.)

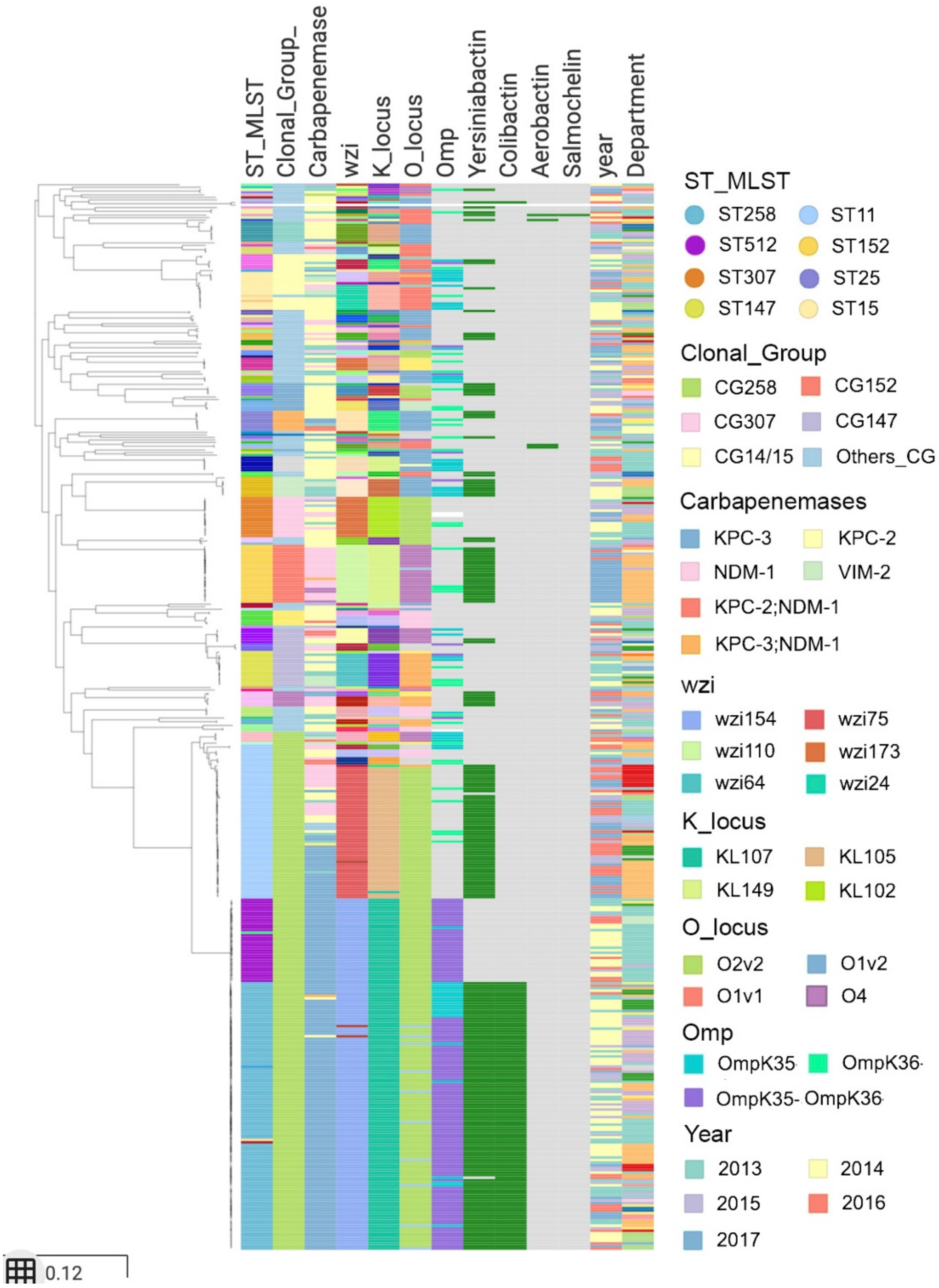


**Supplementary Figure 4.** Microreact visualization of 425 Colombian isolates of carbapenem-resistant *K. pneumoniae*. See visualizations on Microreact at: https://microreact.org/project/vcsgT8Ic4/32d63ab7.
